# On the structure of cortical micro-circuits inferred from small sample sizes

**DOI:** 10.1101/118471

**Authors:** Marina Vegué, Rodrigo Perin, Alex Roxin

**Author notes:** **Corresponding Author**: Alex Roxin. **Author Contributions** M.V. and A.R. did modeling and data analysis and wrote the paper with input from all authors, R.P. collected data.

## Abstract

The structure in cortical micro-circuits deviates from what would be expected in a purely random network, which has been seen as evidence of clustering. To address this issue we sought to reproduce the non-random features of cortical circuits by considering several distinct classes of network topology, including clustered networks, networks with distance-dependent connectivity and those with broad degree distributions. To our surprise we found that all these qualitatively distinct topologies could account equally well for all reported non-random features, despite being easily distinguishable from one another at the network level. This apparent paradox was a consequence of estimating network properties given only small sample sizes. In other words, networks which differ markedly in their global structure can look quite similar locally. This makes inferring network structure from small sample sizes, a necessity given the technical difficulty inherent in simultaneous intracellular recordings, problematic. We found that a network statistic called the sample degree correlation (SDC) overcomes this difficulty. The SDC depends only on parameters which can be reliably estimated given small sample sizes, and is an accurate fingerprint of every topological family. We applied the SDC criterion to data from rat visual and somatosensory cortex and discovered that the connectivity was not consistent with any of these main topological classes. However, we were able to fit the experimental data with a more general network class, of which all previous topologies were special cases. The resulting network topology could be interpreted as a combination of physical spatial dependence and non-spatial, hierarchical clustering.

**Significance Statement:** The connectivity of cortical micro-circuits exhibits features which are inconsistent with a simple random network. Here we show that several classes of network models can account for this non-random structure despite qualitative differences in their global properties. This apparent paradox is a consequence of the small numbers of simultaneously recorded neurons in experiment: when inferred via small sample sizes many networks may be indistinguishable, despite being globally distinct. We develop a connectivity measure which successfully classifies networks even when estimated locally, with a few neurons at a time. We show that data from rat cortex is consistent with a network in which the likelihood of a connection between neurons depends on spatial distance and on non-spatial, asymmetric clustering.

## Introduction

Network architecture shapes the way in which information is transmitted and stored in neuronal circuits. In the mammalian cortex, complex functions such as sensory processing, decision making, memory storage and even abstract thought processes are thought to be the result of a highly structured network topology. Therefore, understanding the structure of cortical micro-circuits may be a key step towards a deep understanding of how the brain performs such tasks.

The organization of cortical micro-circuits varies across brain areas and species, and undergoes continual plastic modifications during the lifetime of a given individual as a result of experience (Trachtenberg et al., 2002; Zuo et al., 2005; Le Bé and Markram, 2006; Hofer et al., 2009). It is accepted, however, that these circuits also exhibit certain regularities, the canonical example of which is a well defined vertical organization into layers. The existence of conserved connectivity principles suggests the notion of a neocortex composed of a juxtaposition of similarly structured building blocks (Szentagothai, 1978; Mountcastle, 1997; Silberberg et al., 2002), which are then dynamically adjusted to respond to the precise demands of every subsystem, in a continuously changing environment.

In the last decades, much effort has been devoted to elucidating the structure of cortical micro-circuits. Intracellular recording techniques have made it possible to assess the presence of monosynaptic connections between pairs of neurons in cortical slices directly (Markram et al., 1997; Mason et al., 1991; Holmgren et al., 2003; Song et al., 2005; Perin et al., 2011). The morphological examination of synaptic contacts via electron microscopy can also in principle provide ground-truth connectivity (Denk and Horstmann, 2004; Bock et al., 2011; Kleinfeld et al., 2011; Kasthuri et al., 2015). Finally, several studies have sought to infer network connectivity from observations of the neuronal dynamics (Nykamp, 2007; Pajevic and Plenz, 2009; Stetter et al., 2012; Sadovsky and MacLean, 2013; Tomm et al., 2014).

One important limitation of cell recording techniques, however, is that they currently allow for the study of only small groups of neurons simultaneously. Therefore, micro-circuitry reconstructions necessarily require an inference process from partial data. Despite these limitations, experimental studies have brought to light some fundamental common principles, such as that the connections tend to be sparse, with connection rates between pyramidal neurons in the range 5-15% (Markram et al., 1997; Le Bé and Markram, 2006; Holmgren et al., 2003; Mason et al., 1991; Ko et al., 2011; Wang et al., 2006). Recent work has also determined specific connection rates depending on the pre- and post-synaptic cell types (Hill et al., 2012; Jiang et al., 2015). Interestingly, there is increasing evidence that the connectivity between pyramidal neurons in different areas and layers is far from the Erdös-Rényi (ER) random network model, where connections appear independently with a fixed probability *p*. These so-called “non-random” features include an excess of reciprocal connections, which can be quantified by the ratio between the number of bidirectional connections and the expected number of such connections in ER networks with equivalent connection rates (*R*). *R* has been reported to be around 2-4 in visual cortex (Song et al., 2005; Wang et al., 2006; Mason et al., 1991), 3-4 in somatosensory cortex (Markram et al., 1997; Le Bé and Markram, 2006) and 4 in mPFC (Song et al., 2005; Wang et al., 2006). Additional evidence for this non-randomness is the over-representation of highly connected motifs (Song et al., 2005; Perin et al., 2011) and the finding that the connection probability between neuron pairs increases with the number of shared neighbors (Perin et al., 2011). Some initiatives are seeking to leverage these data in order to construct realistic micro-circuit models for numerical simulation (Hill et al., 2012; Markram et al., 2015; Reimann et al., 2015; Ramaswamy et al., 2015). On the other hand, a recent theoretical study has shown that some of these features arise naturally in network models that maximize the number of stored memories (Brunel, 2016).

In this paper we have studied several broad classes of network structure that could potentially explain the observed non-randomness. These include clustered networks (Litwin-Kumar and Doiron, 2014), spatially structured networks (Holmgren et al., 2003; Perin et al., 2011; Jiang et al., 2015) and networks defined by strong heterogeneity in the number of incoming and outgoing connections of neurons (Roxin, 2011; Timme et al., 2016).

Surprisingly, all of these network classes were compatible with the reported non-randomness. In fact, we found that networks with qualitatively distinct *global* structure could yield similar statistical features, e.g. motifs, when all the available information came from the study of small groups of neurons, as in experiment. However, we found that a particular combination of motifs, known as the sample degree correlation (SDC), provides a unique fingerprint for each network class, based only on the analysis of small samples of neurons. Using the SDC we showed that micro-circuit data from rat somatosensory cortex (Perin et al., 2011) and from rat visual cortex (Song et al., 2005) were incompatible with any of these network classes. Rather, the data lead us to develop a more general network class which reduces to the previous models under certain constraints. Our results suggest that the non-random features of cortical micro-circuits reflect a combination of spatially-decaying connectivity and additional non-spatial structure which, however, is not simple clustering.

## Materials and Methods

All the networks are treated as directed graphs with *N* neurons. We assume that the network’s size *N* is large and that the network is *sparse*, meaning that its connection density *p* is “small”. We use the following notations: *i → j* a connection exists from neuron *i* to neuron *j*;*i ↦ j* a connection exists from *i* to *j* but not from *j* to *i*; *i ↔ j* there is a bidirectional connection between *i* and *j*.

### Network models

**Erdös-Rényi**(**ER**) Connections are generated independently with probability *p*. ***ER bidirectional***(***ER-Bi***) Connections between a pair of neurons (*i*, *j*) are generated independently according to

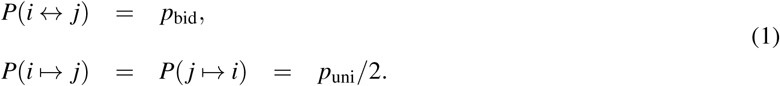

The sparseness and the number of bidirectional connections relative to random are

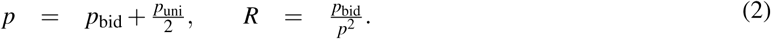

### Clusters (Cl)

Each neuron belongs to one or more clusters and cluster membership is homogeneous across the network. This means that, for any neuron *i*, the number of other neurons that share a cluster with *i* is almost constant. More precisely, if *n_i_* denotes the number of neurons that are at least in one of the clusters of *i*,

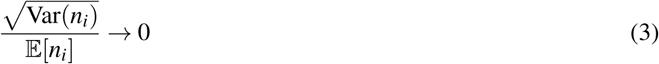
as *N* → ∞.The typical example is a network with a fixed number of clusters *C*≪*N* where each neuron belongs to one cluster that is chosen uniformly at random. In this case, *n_i_* ∼ Binomial(*N* ∓ 1,1/*C*), so

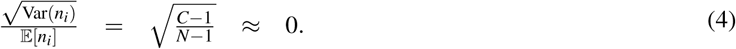

Connections are generated independently with probability *p*_+_ when neurons are in the same cluster and *p*_ otherwise, *p_*< *p*_+_. Defining *f*_+_ and *f_* − = 1−*f*_+_ as the expected fraction of pairs in the same and in different clusters, respectively, *p* and *R* are

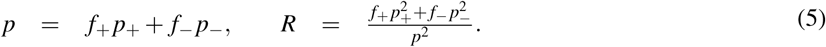

In our simulations each neuron belongs to one cluster which is chosen uniformly at random, so the expected cluster size is *N*/*C* and

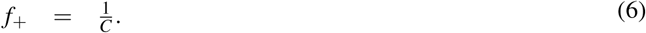

### Clusters with heterogeneous membership (Cl-Het)

Each neuron belongs to zero, one or more clusters but now cluster membership is heterogeneous across neurons, which means that Eq. (3) does not necessarily hold. Connections are defined as in the previous model. In our simulations we have considered networks with *C* ≪ *N* clusters where each neuron has a probability *p_c_* = 1/*C* of belonging to any given cluster. Therefore, neurons can be simultaneously in different clusters and clusters may have non empty overlap. *p* and *R* are given by expression (5) as before, but now the expected fraction of pairs in the same cluster is

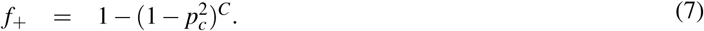

Defining again *n_i_* as the number of neurons that are at least in one of the clusters of *i*,

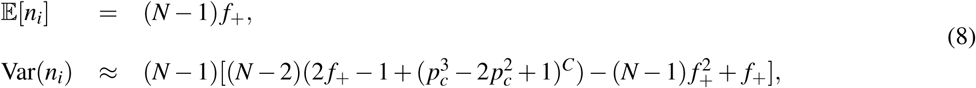
so, if *C* is fixed and *N* is large,

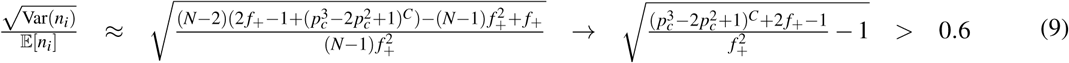
for *C* ≥ 2. This means that there is a non negligible variability across neurons in terms of cluster membership, which has important consequences for the statistics that we will consider later.

### Distance (Dis)

Connections are made independently with a probability that decays with the distance *r_ij_* between the neurons *i* and *j*:

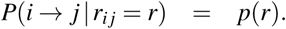

We have

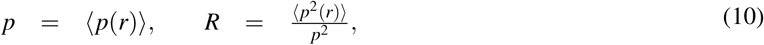
where 〈〉 denotes an average over the distribution of distances in the network. We assume that distances are homogeneously distributed in the network, i.e., that the proportion of neurons that are a given distance away from a neuron *i* does not vary substantially from *i* to *i*. This condition is analogous to requirement (3) for clustered networks. When it does not hold, the model belongs to the Cl-Het class in terms of the properties studied in this paper.

### Degree (Deg)

We consider networks defined by a given joint in/out-degree distribution *f*_(in, out)_ (*k*, *k′*). One realization of the model is obtained by generating a degree sequence 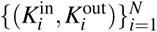 from *N* independent instantiations of *f*_(in, out)_ and uniformly selecting one network among the family of directed graphs that have 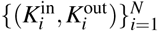 as their degree sequence.

Since the number of edges in any directed network equals the sum of the in-degrees and the sum of the out-degrees, the expectation of the in- and the out-degree have to be equal:

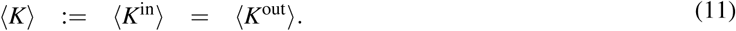

The sparseness is

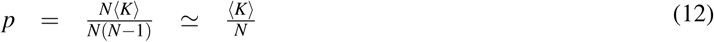
in the large *N* limit.

In this model, the connection probability once the network degrees are known can be approximated by

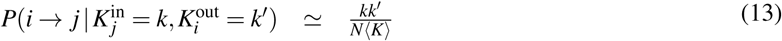
and, since, once conditioned to the degrees of neurons *i* and *j*, *i* →*j* and *j* →*i* can be considered independent events,

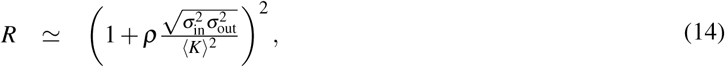
where 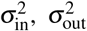 and *ρ* stand for the in/out-degree variances and the Pearson correlation coefficient of individual in/out-degrees, respectively.

### A more general family of networks: the Modulator model

It is possible to consider a very general class of network models in which each neuron *i* has an associated parameter *x_i_* and the connections are made independently with probability

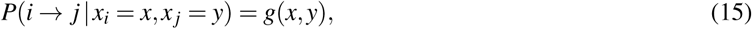
where 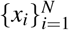 are independent and identically distributed random variables. All the previous models except the ER-Bi can be interpreted, at least locally, as particular cases of this model.

In clustered networks (Cl and Cl-Het), *x_i_* denotes the cluster membership of neuron *i*, whereas in the Dis model, *x_i_* represents the “position” of neuron *i*. In both of these cases the connection probability depends on a notion of distance between pairs, so the function *g* is symmetric: *g*(*x*,*y*) = *g*(*y*,*x*). Moreover, in a random sample of the Cl and Dis models, coexistence in a cluster or distance can be assumed to be independent from pair to pair, as long as the sample size is small compared to the network size. In the Cl-Het model this is not the case by virtue of the neuron-to-neuron heterogeneity in cluster membership: the likelihood of a connection from a neuron *i* is highly dependent on the number of other neurons in the network that share a cluster with *i* (the quantity *n_i_* defined before). Since this quantity varies significantly from neuron to neuron, connections from neuron *i* cannot be assumed to appear independently.

In the Deg model, the connection probability from neuron *i* to neuron *j* once the degrees are known can be approximated by Eq. (13). Additional connections from neuron *i* can be assumed to be made independently as long as *k*≫1. This independence assumption can be extended up to a group of *n* neurons as long as the degrees are large compared to *n* and *n* ≪ *N*. Then, the Deg model becomes a special case of the Modulator model in which x*_i_* = 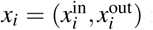 is the 2-dimensional vector of the degrees of *i* and *g*(*x*,*y*) = *g*_1_(x)*g*_2_(*y*), where 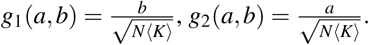

### In/out-degree correlation in small samples

Given a random sample of a network, we define the sample degree correlation (SDC) as the Pearson correlation coefficient between in- and out-degrees of individual neurons in the sample:

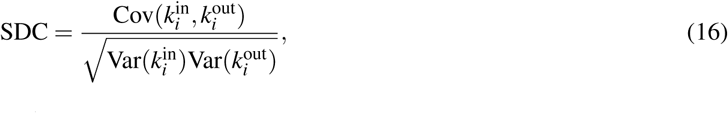
where *i* represents a random neuron and 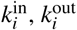 are the in- and out-degrees of *i* in the sample.

In order to compute the SDC in our models we first need to introduce the following statistics. Given any network and random nodes *i*, *j*, *k*, we define

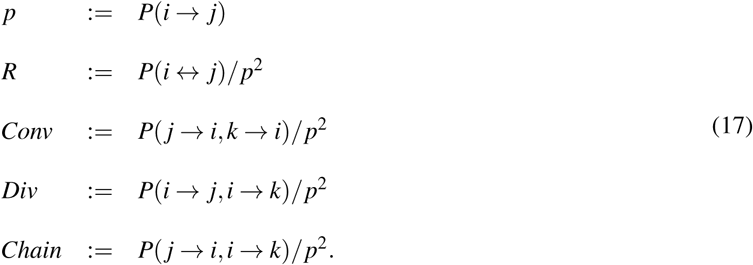

Note that these quantities do not trivially coincide with the motifs first defined in (Song et al., 2005) and reproduced here in Fig.2A. For example, the occurrence of the convergent motif number 5 above chance in Fig.2A can be written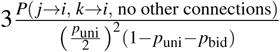, where *p*_uni_ = 2*p*(1 − *pR*), *p*_bid_ = *p***^2^***R* and the factor 3 accounts for the different permutations of *i*, *j* and *k* which produce the same topological configuration. The motifs needed to compute the SDC are not conditioned on the presence or absence of any additional structure in the neuron triplet, merely the existence of, for example, a convergent motif. Therefore, our *Conv* motif is actually a weighted sum of all motifs in Fig.2A containing at least one convergent node, i.e. 5, 7, 9-10, 12-16.

**Figure 2.**
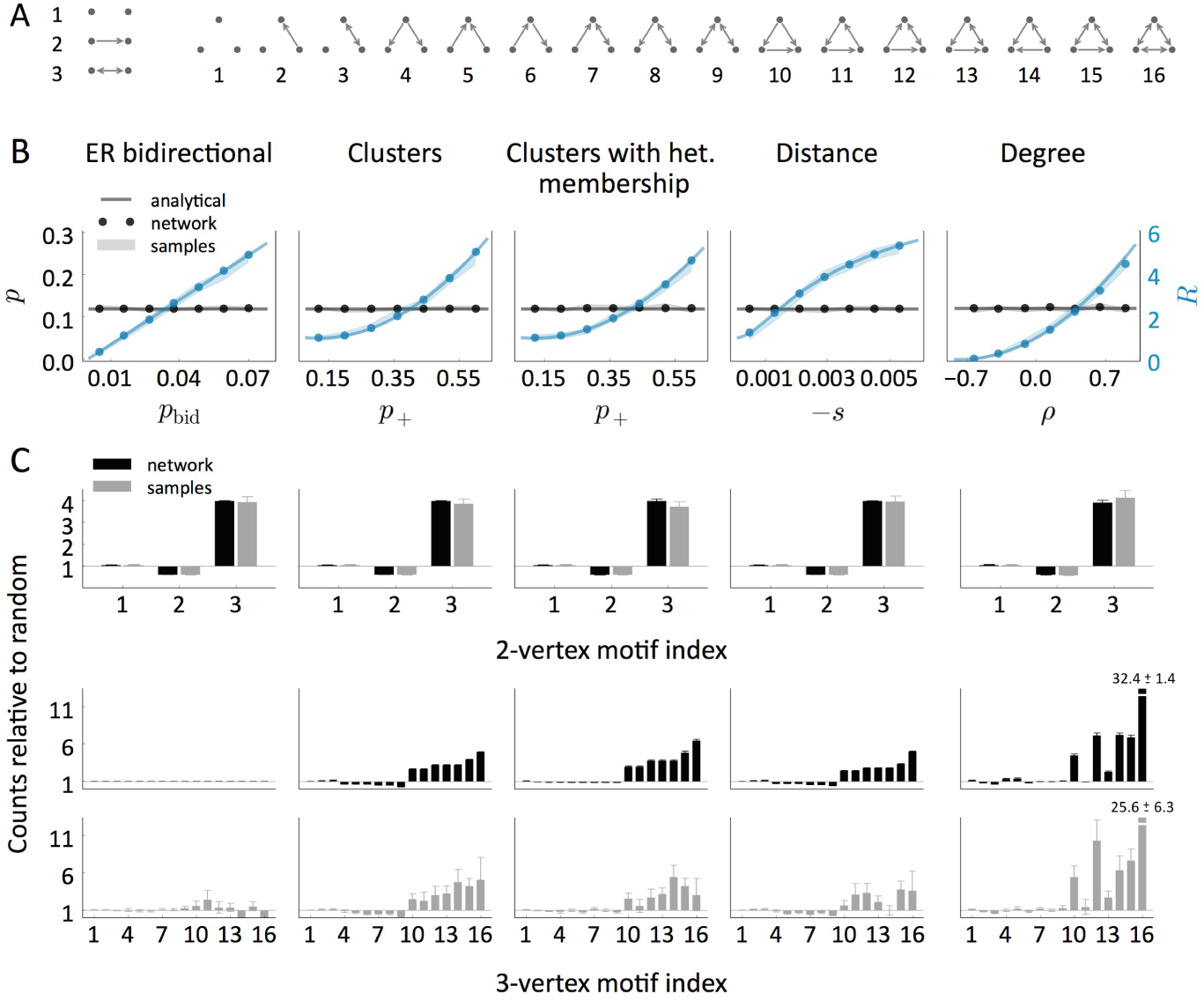
Counts of 2- and 3-neuron motifs relative to random models. **A** Representation of all the possible 2- and 3-neuron motifs. **B** Sparseness (*p*) and expected number of reciprocal connections relative to random (*R*) as a function of a model parameter. In all the models except the Deg, an additional parameter was varied (p_uni_, *p*_, *p*_, *t*, respectively) to keep *p* constant. In the Dis model, neurons are arranged in a ring and the connection probability as a function of distance *r* is defined by the sigmoid function 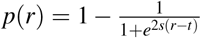, so *t* is the point where the absolute slope is maximal and –s is this absolute slope. **C** Counts of all the 2- and 3-neuron motifs relative to random models (ER and ER-Bi, respectively) in networks with *p* = 0.12, *R* = 4. We used 5 different networks of size *N* = 2000 per condition. The computations were performed both on the whole network and on 163 samples of size 4 per network. Shaded regions and error bars indicate mean ± **SEM.**

The in- and out-degrees of a node *i* in a sample of size *n* can be expressed as

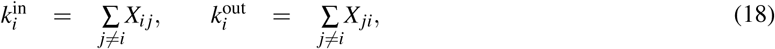
where *X_ij_* = 1 whenever *j → i* and *X_ij_* = 0 otherwise (the sums in (18) are over the *n* indices of the neurons in the sample). Explicitly computing the sample degree variances and the covariance between in- and out-degrees of neuron *i* from expression (18) we find

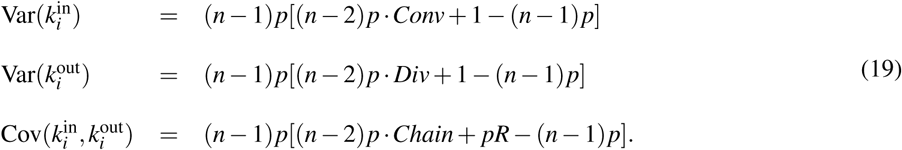

In the ER-Bi model, the pair to pair independence implies that *Conv* = *Div* = *Chain* = 1 and

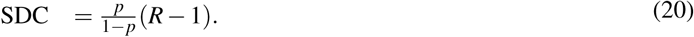

In the Modulator model, the quantities *p*, *R*, *Conv*, *Div*, *Chain* can be rewritten in terms of moments of *g*:

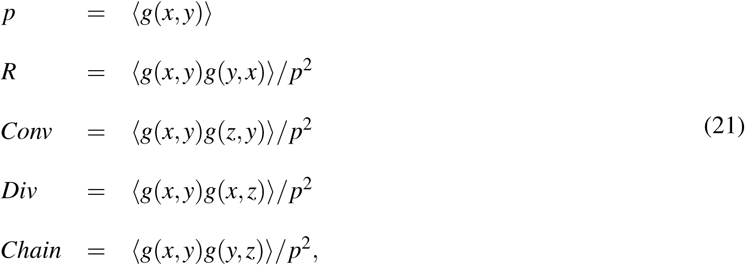
where 〈〉 indicates an average over the distribution of *x*, *y*, *z*, which are independent and identically distributed random variables. We have the following particular cases:

i. If *g*(*x*,*y*) can be assumed to be independent of *g*(*x*,*z*), *g*(*z*,*x*), *g*(*z*,*y*) and *g*(*y*,*z*), then *Conv* = *Div* = *Chain* = 1 and

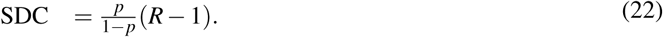 In the Cl and Dis models, the property of being in the same cluster (Cl) and the distance between a pair (Dis) can be assumed to be independent from one pair to another, so (22) is a good approximation of the sample degree correlation.
ii. If *g* is symmetric, that is, *g*(*x*, *y*) = *g*(*y*, *x*), then *Conv* = *Div* = *Chain* and

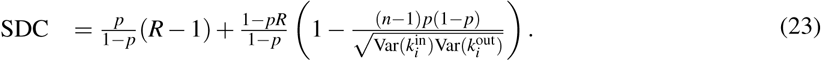 This is the case of the Cl-Het model. Note that in the Cl/Dis models *g* is also symmetric, so this expression for SDC is a generalization of (22), which is recovered whenever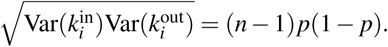
iii. If *g* is multiplicative, that is, *g*(*x*, *y*) = *g*_1_ (*x*)*g*_2_(*y*), then *Chain*^2^ = *R* and

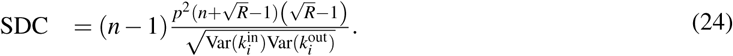 The Degree model fits within this case.

### Generation of distance-dependent networks

In the simulations of Figs. 1 to 6A we considered neurons arranged in periodic rings where *r* € {0,1, ···, [*N*/2]} and

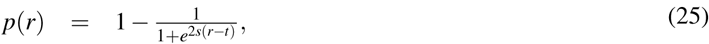
which defines a decreasing sigmoid function whose absolute slope is maximal at *r* = *t* and its value is *-s.* In the simulations of Fig. 6C,D we also included two-dimensional periodic lattices where *r* ϵ 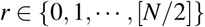 and *p*(*r*) was given by Eq. (25).

**Figure 6.**
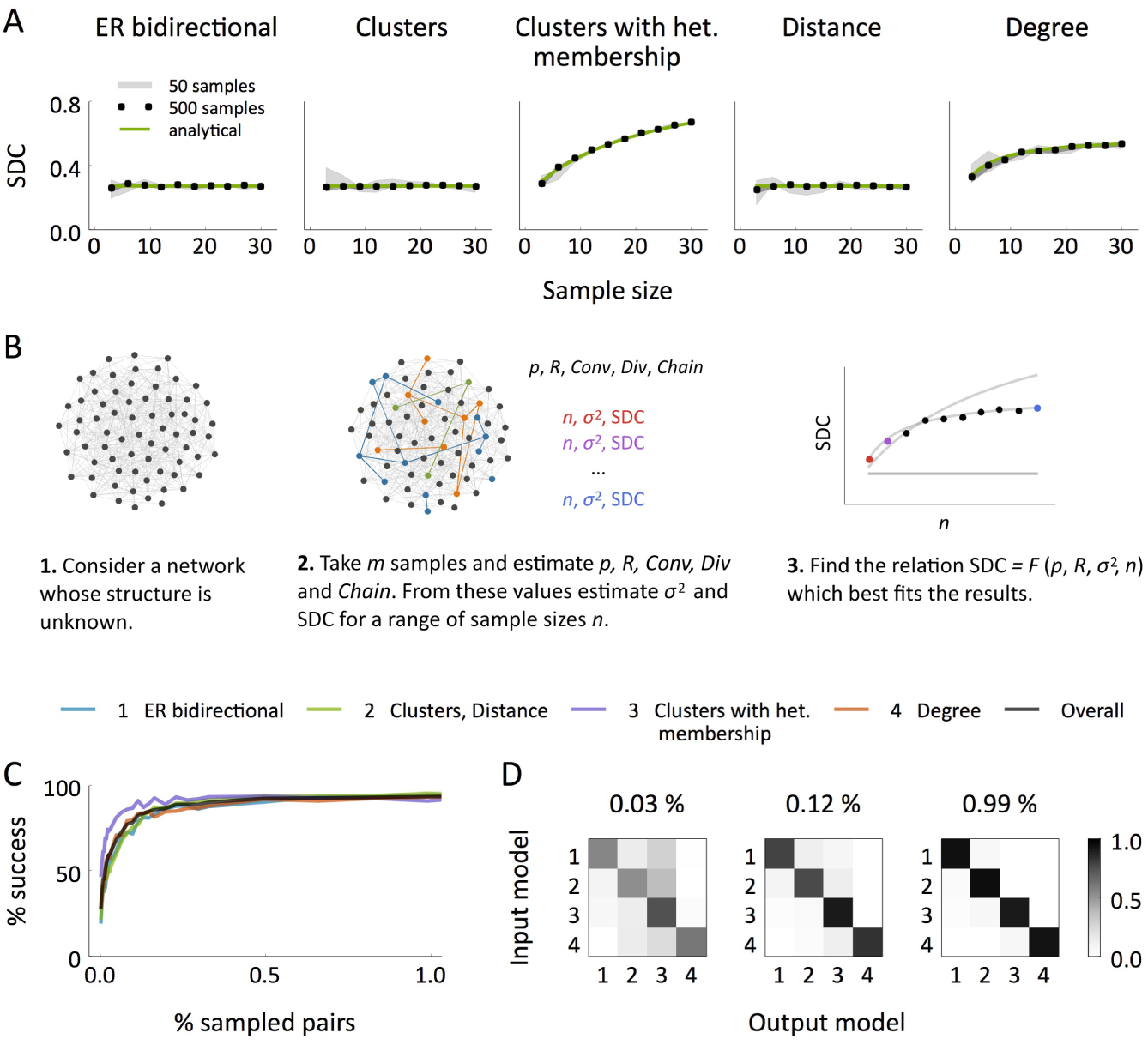
Sample in/out-degree correlation (SDC) as a measure to distinguish between classes of networks. **A** SDC in samples of 3 to 30 neurons in the different models. In all the networks, *N* = 2000, *p* = 0.12, *R* = 3. We computed the empirical correlations using 50 and 500 samples per network for each sample size. Every analysis was performed independently in 5 different networks and the shaded region indicates the resulting mean ± SEM. **B** Schematic representation of the algorithm proposed to distinguish between the model classes: (1) ER-Bi, (2) Cl/Dis, (3) Cl-Het and (4) Deg. **C** Success rate of the algorithm performed on randomly generated networks with *N* = 2000, *p* ∈ [0.05,0.23], *R* ∈ [1.5,4.1] as a function of the percentage of neuronal pairs analyzed. Each success rate was computed over 2000 experiments. **D** Frequencies of all the possible input-output combinations in the experiments shown in C, for three choices of the proportion of analyzed pairs. Each frequency is normalized by the frequency of the input model so that the sum of every row is 1.previously reported in Perin et al. (2011), these data show a clear dependency of connection probability on intersomatic distance. The estimated connection density and number of reciprocal connections relative to random were *p* = 0.144, *R* = 2.575. The analysis of the SDC revealed a relationship which deviates from any of the previously defined models, Fig. 7A. Although the form of the SDC appears close to that of the Dis model (Fig. 7A left), the degree variance from the data *σ*^2^, which should be that of a Binomial distribution, differs strongly from the theoretical value (Fig. 7A right). Note that the degree variance for the other two classes of network is a free parameter and hence here is estimated directly from the data.

### Generation of networks from a prescribed in/out-degree distribution

To generate networks according to the Deg model we have used the following method: given a joint distribution defined by 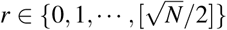we independently assign to each node *i* a pair 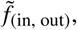. Then we create each connection *i*→ *j* independently with probability 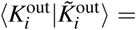. The final degrees in the network satisfy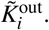.Despite the resulting degree distribution in the network is no longer given by 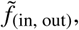 the statistics *〈K〉* and Cov*(K^in,^K^out^)* are preserved (assuming that *N* is large and 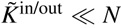. The degree variances become larger, in particular 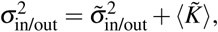, and this results in the correlation coefficient being smaller, 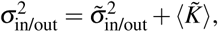.

In all our simulations, the variables 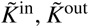 followed Gamma distributions with a shift of magnitude *D* > 0. In almost all our simulations they had to be positively correlated and we defined them in the following way: if *X* ∼ Gamma(*κ*_1_, θ) and *Y*, *Z* ∼ Gamma(*κ*_2_, *θ*) (*κ*, *θ* > 0) are independent random variables, we set

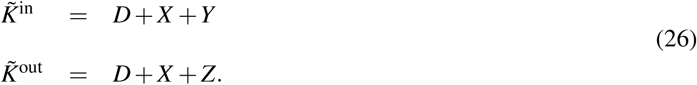

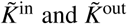 follow *D*-shifted Gamma(κ = κ_1_ + κ_2_,θ) distributions and their correlation coefficient is 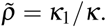. In Fig. 2B we also constructed networks with negative degree correlation. In this case we first generated 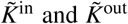 independently and then we inversely ordered the two sequences 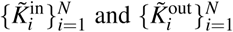 By reordering a fraction of values in one of the two sequences we could adjust the correlation coefficient.

### Implementation of the SDC criterion on a random network generator

In Fig. 6C,D we applied the SDC criterion on networks generated randomly according to the models ER-Bi, Cl/Dis, Cl-Het and Deg. We chose a network class and values for *P* ∈ [0.05, 0.23] and *R* ∈ [1.5,4.1] uniformly at random. In the ER-Bi model these parameters determine *p*_uni_ and *p*_bid_. If the chosen class was Cl/Dis, we chose one of these two models with equal probability. In the Cl case, we selected the number of clusters randomly and then computed *p_+_* and *p_−_* to get the desired *p* and *R*. In the Dis case, we chose a dimension (1 or 2) randomly and then placed neurons in periodic lattices of the given dimension. Then we determined the parameters *s* and *t* of Eq. (25) to fit *p* and *R.* If the selected model was Cl-Het we did exactly the same as in the Cl case. Finally, in the Deg model we chose *D* and ρ > 0 randomly and then found θ, κ_1_ and κ_2_ to fit *p* and *R*.

To classify a network according to the SDC, we took m random samples of size *n′* = 12 each. From them we estimated *p, R, Conv, Div* and *Chain* (Eq. (17)) and computed the connection probability as a function of the number of common neighbors. From *p, R, Conv*, *Div* and *Chain* we predicted 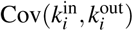for any sample size *n* through Eq. (19). We compared the resulting SDC (seen as a function of n) with the SDC that would result in each of the model classes given the observed *p*, *R* and *σ*^2^ (Eqs. (22), (23) and (24)). We determined which of these relationships between SDC and *n* better described the results by computing the sum of the squared distances between the actual SDC and the model predictions while varying *n*. The range of *n* values used to make this comparison is arbitrary. We chose *n* ∈ {3, ···, 12} but the results are essentially the same for other choices. Since the formula for the Cl-Het model generalizes the formula for ER-Bi/Cl/Dis, the SDC of a network of the class ER-Bi/Cl/Dis will be fitted equally well by these two formulas. Thus, whenever the best fit corresponded to the Cl-Het class, we further studied if the SDC increased significantly with *n* by computing the slope of its linear regression and deciding if it was larger than a critical value *s*,* which had been previously determined by means of simulations. If the slope was smaller than *s\**, the network was reclassified as ER-Bi/Cl/Dis. Finally, to distinguish between ER-Bi and Cl/Dis networks, we determined if the connection probability in the *n′* samples increased significantly with the number of common neighbors. Again, this was done by computing a linear regression and comparing the slope with a previously defined threshold.

We further checked that the same algorithm works if *σ*^2^ and 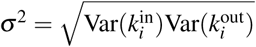 are calculated directly for each *n* on *n*-neuron samples instead of being estimated from *p, R, Conv, Div* and *Chain.* The *n*-neuron samples in this case are subsamples of the original samples of size *n′*. The only limitation of this procedure is that the original sample size *n′* has to be large enough to make it possible to compute *σ*^2^ for *n* in the desired range, whereas the estimation of *p, R, Conv, Div* and *Chain* only requires 3-neuron samples. A study based on sampling from triplets or quadruplets, however, would not allow us to distinguish between the ER-Bi and Cl/Dis classes using the common neighbor rule.

### Implementation of the SDC criterion on data

To apply the SDC criterion on the experimental data we considered all the possible subsamples of the original samples. For each subsample size, we considered in- and out-degrees of all the neurons to compute *σ*^2^ and 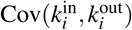 We estimated the data SDC, the predicted SDC for the model classes and their standard errors by means of the Bootstrap method with 1000 re-samplings. We repeated the same procedure considering the predicted *σ*^2^ and 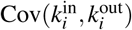 from *p, R, Conv, Div* and *Chain.* The results are almost identical.

### Definition of the model that fits the data

In the proposed model to fit the data of Perin et al. (2011), connections are created independently with probability 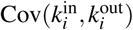. The distance dependency has the form

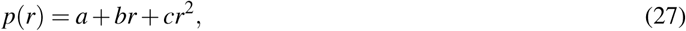
where *r* is the normalized distance 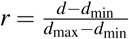∈ [0, 1] that is computed from the real distance *d* in *μ*m and minimal and maximal distances derived from the data, *d*_min_ = 10*μ*m, *d*_max_ = 350*μ*m. We took *a* = 1, *b* = -1.04, *c* = 0.21. The modulatory part is

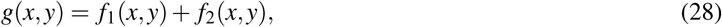
where *f_1_* and *f*_2_ have the form

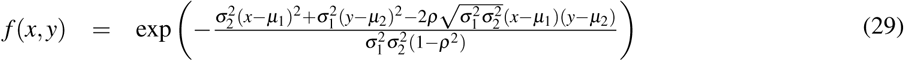
and their parameters are shown in Table 1. The modulators {*x_i_*}*_i_* are independent from neuron to neuron and are drawn from a Gaussian distribution with mean 0 and standard deviation 0.5.

**Table 1.** Parameters of the modulatory function

| | $\mu_1$ | $\mu_2$ | $\sigma_1^2$ | $\sigma_2^2$ | $\rho$ |
| --- | --- | --- | --- | --- | --- |
| $f_1$ | 0.0 | 0.0 | 0.3 | 0.3 | 0.92 |
| $f_2$ | -0.5 | 0.5 | 0.07 | 0.07 | -0.62 |

To obtain a distribution of distances in the simulated data close to the sampled distances in the experiment, we directly generated samples as in the real experiment. In each sample, the first neuron was located in the origin of coordinates and the others were sequentially located on the same plane at a position obtained by drawing a random angle α∈[0,2π] and a radius *r* from a Gamma(*κ*, *θ*) distribution, κ = 3.26, θ = 0.08. The radius was then rescaled as *d* = *d*_0_ + (*d*_1_ - *d*_0_) * *r*, *d*_0_ = 16*μ*m, d_1_ = 250*μ*m. We avoided having neurons too close in space by checking, at every step, if the last neuron was closer than a limit distance *d*_lim_ = 14μm to the already created neurons in the sample. In this case we chose a new position.

## Results

### Canonical network models for cortical circuits

We asked ourselves to what extent simple, canonical models of network topology could reproduce the salient statistics from actual cortical circuits in slice experiments. The simplest possible sparsely connected network model is the so-called Erdös-Rényi (ER) network, for which connections between neurons are made with a fixed probability *p*. However, data show that cortical circuits are not well described by the ER model, and in particular, the occurrence of certain cortical motifs is above what would be expected from ER. Therefore, we consider other candidate network models which go beyond ER (see Fig. 1): i. An ER network with additional bidirectional connections (ER-Bi). This model has just two parameters: the probability of a unidirectional connection p_uni_ and that of a bidirectional connection *p*_bid_. ii. A network with C clusters where cluster membership is homogeneous across neurons (Cl). The probability of connection between neurons within the same cluster is p_+_ while between clusters it is *p*_*_< *p*_+_. iii. A network with C clusters and heterogeneous membership (Cl-Het), where neurons belong to a variable number of clusters. The probability of connection within and between clusters is as for the Cl model. iv. A network with distance-dependent connectivity (Dis), i.e. the probability of connection between two cells at a distance *r* is *p*(*r*), which is a decreasing function of *r*, and v. A network defined by the distribution of in-degrees and out-degrees (Deg), with mean degree *〈K〉*variances 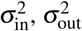 and degree correlation *ρ* (see Materials and Methods for details).

**Figure 1.**
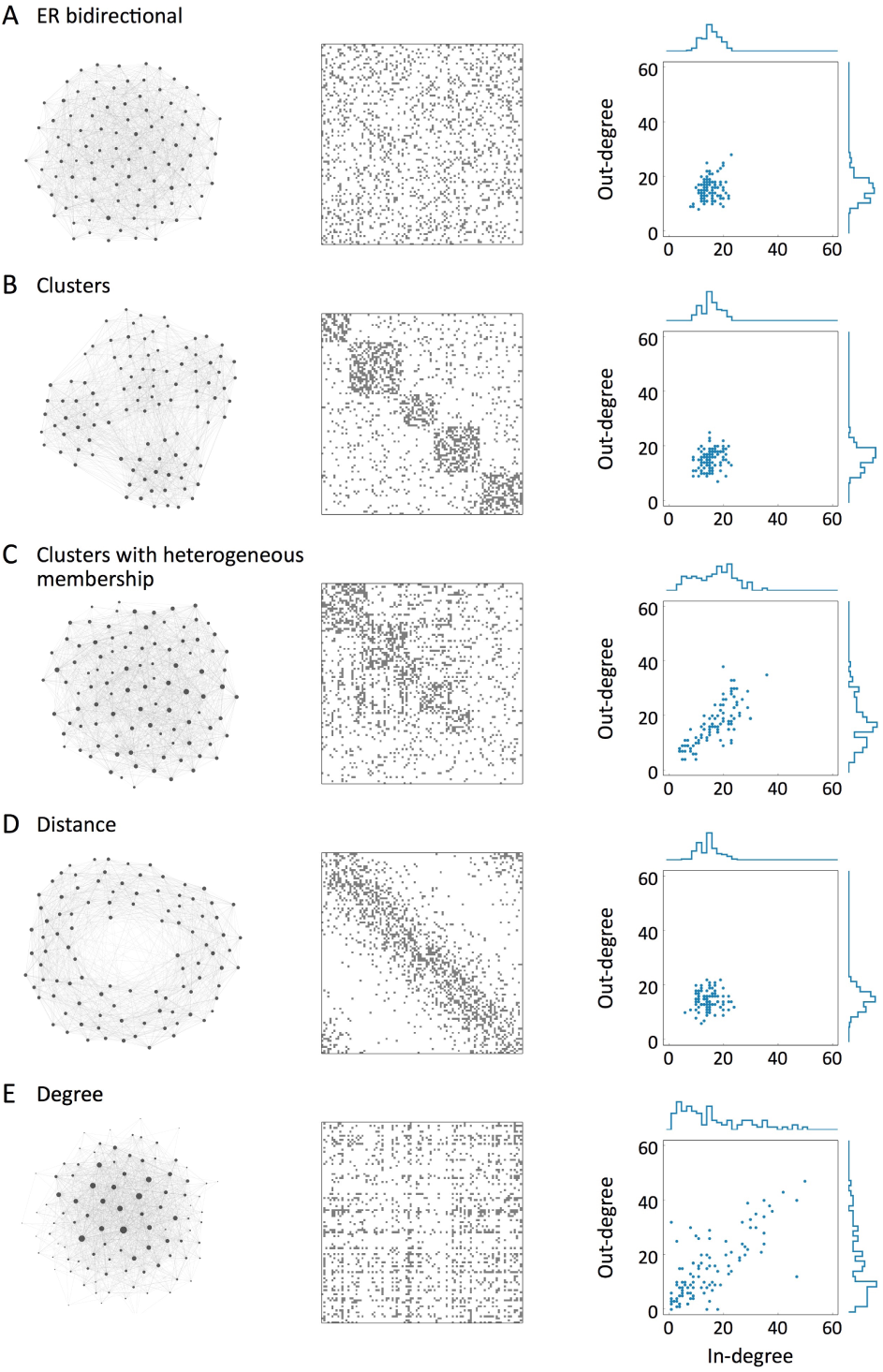
Schematic representation of the models: connectivity (left), adjacency matrix (middle) and in/out-degree distribution (right). The nodes in the left column are arranged according to the ForceAtlas algorithm using Gephi software (Bastian et al., 2009). The size of each node is proportional to the sum of its degrees and the direction of the connections has been omitted for simplicity. In all the networks, *N* = 100, *p* = 0.15, *R* = 2.

### Representation of 2- and 3-vertex motifs relative to random

We first asked whether the deviation in the number of two-neuron motifs relative to random that has been reported previously (e.g. Song et al. (2005)), could be explained by any of the models presented here. Given the sparseness *p* of a network model (that is, the expected number of connections divided by the total number of possible connections), we denote by *R* the expected number of reciprocal connections relative to that in ER(*p*), which can be calculated for each model as shown in Table 2 (see Materials and Methods for details). The expected number of uni-directionally connected and unconnected pairs is then uniquely determined once *p* and *R* are known.

**Table 2.** Sparseness (*p*) and fraction of bidirectional connections relative to random (*R*) in the different models

| Model | $p$ | $R$ |
| --- | --- | --- |
| ER bidirectional | $p_{\text{bid}} + \frac{p_{\text{uni}}}{2}$ | $\frac{1}{p^2} p_{\text{bid}}$ |
| Clusters | $f_+ p_+ + f_- p_-$ | $\frac{1}{p^2} (f_+ p_+^2 + f_- p_-^2)$ |
| Distance | $\langle p(r) \rangle$ | $\frac{1}{p^2} \langle p^2(r) \rangle$ |
| Degree | $\frac{\langle K \rangle}{N}$ | $\left( 1 + \rho \frac{\sqrt{\sigma_{\text{in}}^2 \sigma_{\text{out}}^2}}{\langle K \rangle^2} \right)^2$ |

Once *p* has been fixed, all models can account for a wide range of values in *R*, including the specific values reported in Song et al. (2005); Wang et al. (2006); Mason et al. (1991); Markram et al. (1997); Le Bé and Markram (2006), see Figs. 2B and C (in Fig. 2C we have taken the values of *p* and *R* reported in Song et al. (2005)). The numbers of three-neuron motifs relative to ER-Bi are also qualitatively similar across models, and consistent with experiment, with the exception of ER-Bi which has no additional structure beyond two-neuron motifs (Fig. 2C, bottom).

**Table 3.** In the models with clusters, *f_* = 1 – f+ and f+ is the fraction of neuronal pairs that are in the same cluster. The brackets 〈〉 in the Dis model represent averages over the distribution of distances in the network. See the main text for a description of the other parameters.

An important question to be addressed here is to what extent the experimental results are sensitive to the sampling procedure. Data are collected through simultaneous patch-clamp recordings and hence can only record from a small number of cells at a time. The motif counts are local properties whose *averages* do not depend on the sample size, but the results can be highly variable if the number of samples studied is not large enough. In order to mimic the experiment by Song et al. (2005), we computed *p* and *R* not only from the study of the whole network but also through 163 samples of 4 neurons per network over 5 networks. As shown in Figs. 2B and C (grey bars), the estimates of the 2-neuron motif counts are quite close to the real counts in networks of *N* = 2000 neurons, which suggests that the magnitudes *p* and *R* are well approximated even when only a small fraction of the total network is known. Although the results of 3-neurons motifs were roughly consistent between the full analysis and that from small sample sizes, they were much more variable than the 2-neuron motifs.

Nonetheless, at least in the example networks shown in Fig.2C, it seems that the particular distribution of triplet motifs might provide a means of classifying the different models. In subsequent sections we will show that there is a particular combination of dual and triplet motifs from which we can extract information about the network class, independently of the choice of other parameters.

### Connection probability as a function of the number of common neighbors

A common neighbor to neurons *i* and *j* is a third neuron which is connected to both *i* and *j*. Perin et al. (2011) have shown that the probability of connection between pairs of cortical neurons increases with the number of common neighbors they have (the so-called “neighbor rule”). Fig. 3 (top) shows the connection probability as a function of common neighbors for examples from each model class from the analysis of a network of 2000 neurons where *p* and *R* are close to the values reported in Perin et al. (2011). In the ER-Bi model, as in the classical ER model, all the pairs are connected independently and according to the same rule, so the number of common neighbors does not provide any information about the “laws” controlling a given connection. All the other models, however, exhibit the common neighbor rule for a general choice of the network parameters. Interestingly, the precise shape of this dependence is quite distinct for different models, indicating it might provide a signature for inferring the full network structure from this one measure. However, these qualitative differences between models largely vanish when realistic sample sizes are analyzed, Fig. 3 (bottom). It is important to keep in mind that the curves shown in Fig.3 are for a particular choice of network from each model class. The exact shape of the curves will depend on that choice. In general, we can say that given small sample sizes one will observe a monotonically increasing dependence of the connection probability on the number of common neighbors for all models but ER-Bi. Specifically, for clustered (distance dependent) models, neuron pairs with more common neighbors are more likely to belong to the same cluster (be closer together), which increases the probability of connection. In the Degree model neuron pairs with more common neighbors are more likely to have large degrees, which again increases the probability of connection.

**Figure 3.**
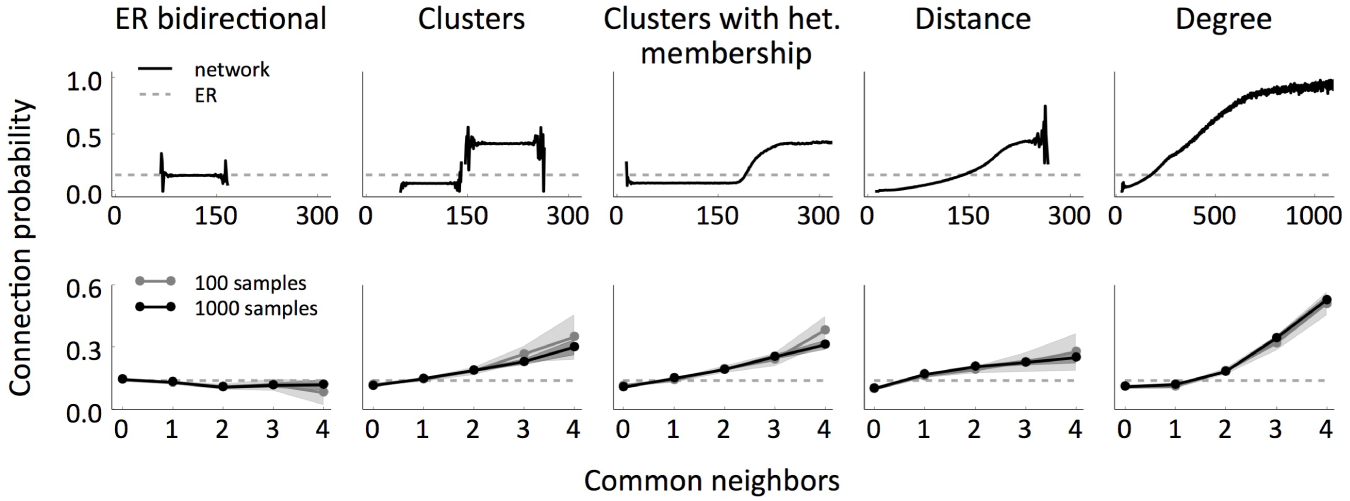
Connection probability as a function of the number of common neighbors for the different models, in the whole network (top) and in samples of size 12 (bottom). In all the cases, *N* = 2000, *p* = 0.14, *R* = 2. The analyses were performed on 5 networks and the shaded regions indicate the resulting mean ± **SEM**. In the sample analyses we took 20 samples per network (100 in total, grey) and 200 samples per network (1000 in total, black). The dotted lines show the expected probability if it were independent of the number of common neighbors, as in the ER and ER-Bi models.

### Degree distributions and higher-order connectivity

Fig. 4A (top) shows the in-degree distributions exhibited by example networks from the different models for physiological values of *p* and *R.* For both the Cl-Het and Deg models the distribution differs dramatically from that of the equivalent ER network. Nonetheless, and as was the case with the common-neighbor rule, when the distributions are constructed from realistic sample sizes (here 12), all models are qualitatively similar, see Fig. 4A (bottom). In fact, due to additional degrees of freedom that both the Cl-Het and the Deg models have, it is possible to define networks with a fixed *p* and *R* whose distributions are nevertheless very different (Fig. 4B). In some situations, the distribution is quite close to ER/ER-Bi cases.

**Figure 4.**
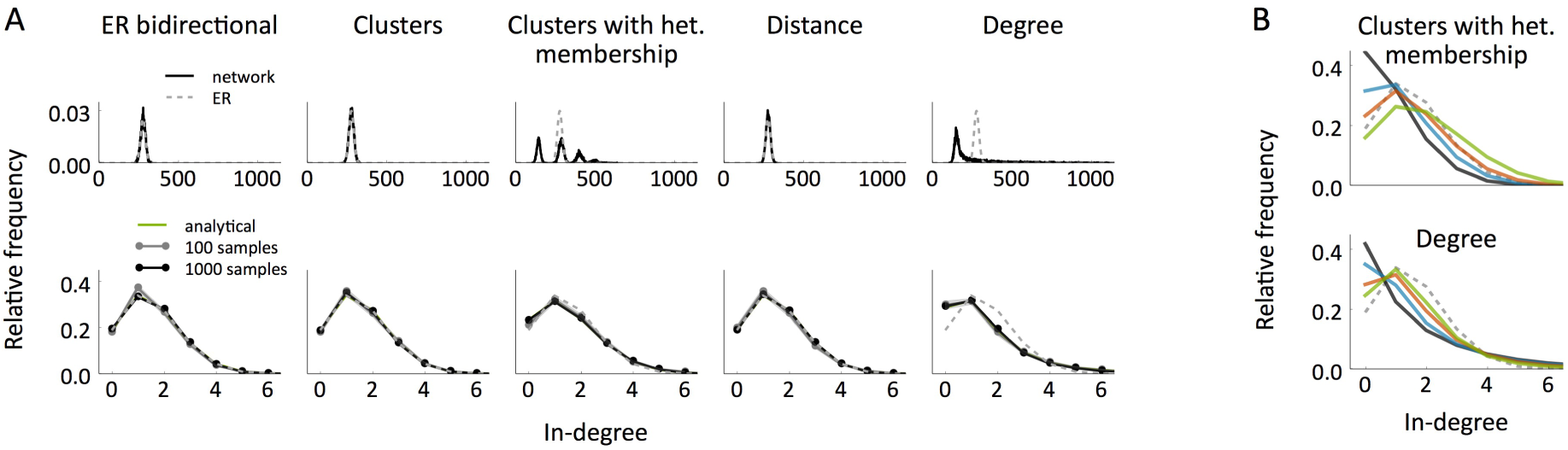
In-degree distribution of the different network models. A In-degree distribution in the whole network (top) versus in-degree distribution in samples of size 12 (bottom) and comparison with the distributions exhibited by the ER model (dotted lines). The networks and samples used are the same as in Fig. 3. The shaded regions indicate mean ± **SEM. B** In-degree distributions in samples of size 12 for different networks generated according to the Cl-Het (top) and the Deg (bottom) models, all of them with *N* = 2000, *p* = 0.14, *R* = 2. network when only local information is available. We found such a measure in the *sample in/out-*degree correlation

Finally, real data also exhibit a significant over-representation of densely connected groups (Perin et al., 2011). We therefore also studied the distribution of the number of connections in small groups of neurons and found that all models, with the exception of ER-Bi, could account for these findings, see Fig. 5.

**Figure 5.**
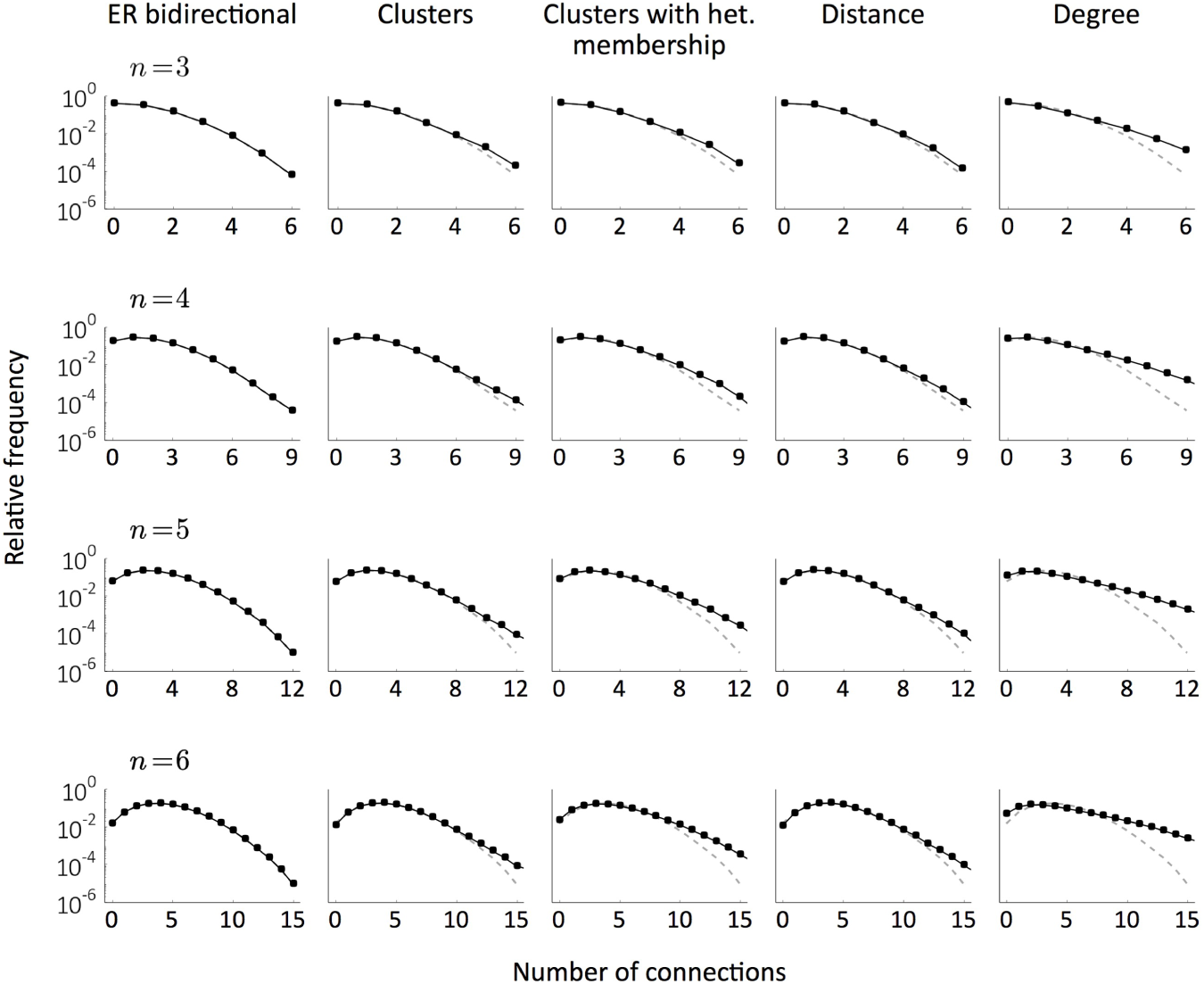
Distribution of the total number of connections in samples of sizes *n* ∈ {3,…, 6} for the different models (black) compared to the distribution obtained in ER bidirectional networks with the same *p* and *R* (dashed grey). The parameters are the same as in Figs. 3 and 4. The analyses were performed on 5 networks per condition and the computations come from 10^5^ random samples for each network.

### A method for distinguishing between network models using measures from small sample sizes

We sought a measure, based on small sample sizes, which would allow us to distinguish between the classes of topological models defined here. In other words, we looked for a way to infer general topological properties of the

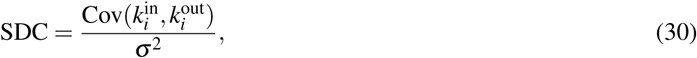
where 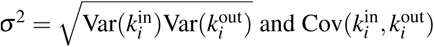and *i* represents a random neuron in the sample. The SDC therefore depends on the variances and covariances of the sample degrees. The in-(out-)variance in turn depends on the occurrence of convergent (divergent) motifs, while the covariance depends on the occurrence of chain and reciprocal motifs. All of these quantities can be calculated analytically for the network classes we have considered here, and the SDC is finally expressed as a function of *p*, *R*, *σ*^2^ and the sample size *n*, see Materials and Methods for details. In particular, we can group the five network types into three classes based on the functional form of the SDC: (1) For the ER-Bi, Cl, and Dis models we find SDC = 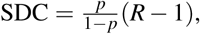 For these models the sample degrees follow distributions which are very close to Binomial(*n* – 1, *p*) as in the classic ER model; in the ER-Bi model by definition and in the other two cases because the distance and the property of belonging to the same cluster are almost independent from pair to pair in a small sample. (2) For the Cl-Het model 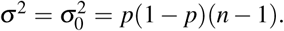 and (3) for the Deg model SDC =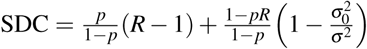 The SDC is therefore independent of the sample size *n* (as long as it is small compared to the network size, *n/N* ≪1) for the first class of networks, while for the Cl-Het and Deg models it changes with *n* in functionally different ways (Fig. 6A). We can additionally use the common-neighbor rule to distinguish between the ER-Bi (which shows no dependence) and the Cl and Dis models (which do). Note that these four classes of networks have SDC ≡ 0 whenever *R* = 1, which means that networks that do not show an over-representation of bidirectional connections cannot be distinguished in terms of the SDC. Therefore, as long as *R* > 1, in principle we can distinguish between all models, except for the Cl and Dis models. This is not surprising given that the Cl is nothing but a particular case of the Dis where the distance is binary.

We applied this “SDC criterion” to networks of size *N* = 2000 generated randomly according to the four classes of models presented here (grouping Cl and Dis), with *p* and *R* chosen uniformly in the ranges [0.05,0.23] and [1.5,4.1], respectively. We then used the SDC to distinguish between the different model classes by taking samples of size *n′* = 12. We estimated *σ*^2^ and the SDC over a range of sample sizes *n* by computing *p*, R and the occurrence of divergent and chain motifs (through the quantities *Conv*, *Div* and *Chain* defined in Eq. (17)), see Fig. 6B and Materials and Methods for details. An alternative approach is to generate n-neuron subsamples from the original samples of size *n′* and directly compute *σ*^2^ and the SDC for each *n* ≤ n′. We also checked that the performance is almost the same for the second method when *n′* = 12 (data not shown). The advantage of estimating *σ*^2^ and the SDC instead of calculating them directly is that it allows to implement the criterion even when the original samples are small (e.g. *n′* = 3,4). To further distinguish between the ER-Bi and Cl/Dis classes we studied if the connection probability increases with the number of common neighbors in the *n′* samples.

The efficacy of this classification criterion increases with the number of samples considered, *m*. Fig. 6C,D shows the performance as a function of the percentage of neuronal pairs analyzed. The rate of success is above the chance level (which is 25%) for all models already for *m* = 2 samples (0.007% of pairs) and reaches 94% when the proportion of analyzed pairs is about 1%.

Analysis of the SDC in data from rat somatosensory cortex

We implemented our SDC criterion in the data obtained by Perin et al. (2011) from pyramidal neurons of the rat somatosensory cortex. The data come from 6, 9, 5, 10 and 10 groups of 8, 9, 10, 11 and 12 neurons, respectively. As

Since the SDC can be extrapolated when the counts of two- and three-neuron motifs are known, we calculated the expected SDC in putative samples of 3 to 12 neurons from the motif distribution described in Song et al. (2005) (Fig. 7B), which corresponds to layer 5 pyramidal neurons in rat visual cortex. The connection density and the number of reciprocal connections relative to random in this case are *p* = 0.116 and *R* = 4. The results are qualitatively similar to the ones computed directly from the data of Perin et al. (2011). This suggests the underlying network structure itself may be similar.

**Figure 7.**
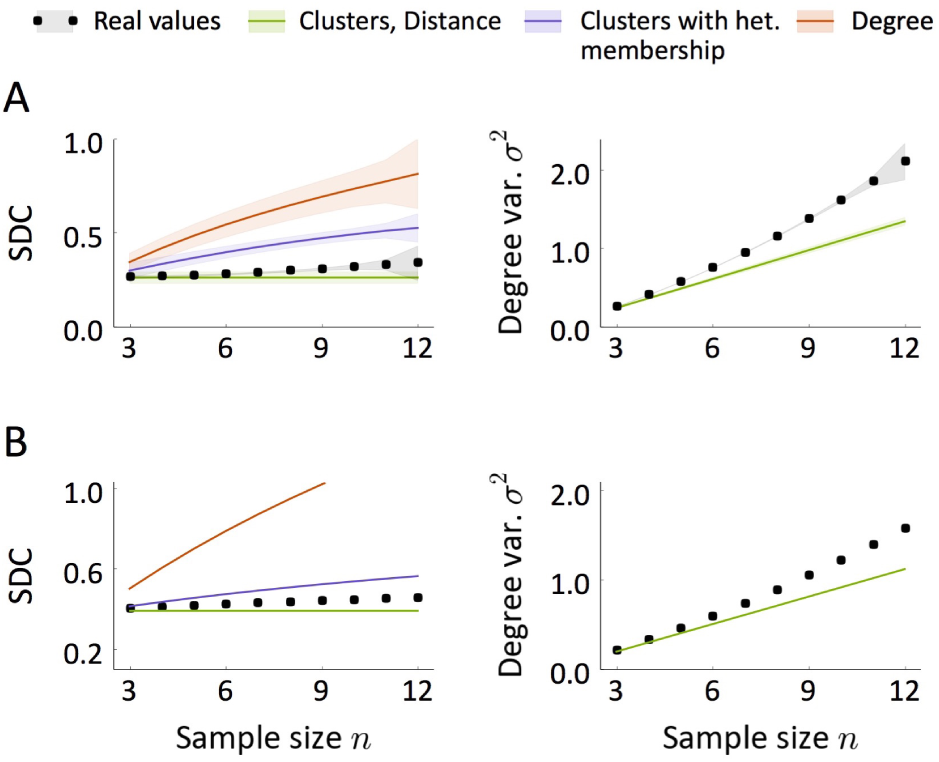
Sample in/out-degree correlation SDC and geometric mean of the sample degree variances *σ*^2^ as a function of the sample size *n.* **A** Values calculated directly from the data of Perin et al. (2011). **B** Inferred values from the motif counts presented in Song et al. (2005). The black curves correspond to the observed SDC and *σ*^2^, whereas colors show the expected SDC (σ^2^) in networks generated according to the studied models with the same p, R, *σ*^2^ (p, R) as in data. Shaded regions in **A** indicate mean ± SEM computed with the Bootstrap method.

### A general class of network model

We discovered that all of the models, with the exception of the ER-Bi model, which could be rejected already by its failure to capture triplet motifs and the neighbor rule, belong to a more general class of model. Specifically, in what we dub *Modulator* networks, the probability of a connection from neuron *i* to neuron *j* is 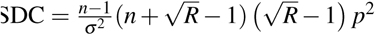where *x_i_* and *x_j_*, the *modulators*, are properties associated with neurons *i* and *j*. The models we have considered so far are special cases of this more general modulator framework.

In the clustered and distance-dependent models that we have considered, *g*(*x*, *y*) = *g*(*y*,*x*) is reflection symmetric. In this case the modulators are the position or membership in a cluster (or group of clusters). It can be shown that any Modulator network with a symmetric *g* exhibits the same SDC as the Cl-Het model. If, additionally, *g*(*x*, *y*) can be assumed to be independent from one neuronal pair to another (as in our Cl and Dis models when a small sample is considered), the formula reduces to the Cl/Dis case. In the Deg model *g* is separable, i.e, *g*(*x*, *y*) = *g_1_* (*x*)*g2*(*y*), and the modulator itself is the pair of in- and out-degrees. The *g* function is just the product of the pre-synaptic out-degree and post-synaptic in-degree, normalized by the appropriate factor. In Materials and Methods we show that the SDC of any Modulator network with separable *g* has the form of the SDC of the Deg model.

Therefore, the SDC criterion not only makes it possible to distinguish between the families Cl/Dis, Cl-Het and Deg, but allows for a classification into three major types of Modulator networks, defined by different properties and symmetries. The fact that the data are not fit by any of the models indicates that real cortical circuits have features which violate the reflection symmetry and separability of the function g.

Since the estimated SDC lies in between the predicted SDC for the Dis/Cl and Cl-Het models (Fig. 7A), one would be tempted to think that a hybrid network from these two classes would be compatible with data. Such a model, however, would still belong to the class of Modulator networks with symmetric *g* and would therefore exhibit the same SDC as the Cl-Het class (purple line in Fig. 7A). This suggests that not only is there additional structure in the data beyond the distance dependence of connection probabilities, but that this structure is not simple clustering.

### Data are consistent with network with spatial dependence and hierarchical clustering

We were able to obtain an excellent fit to all relevant topological statistics in the data with a Modulator network. Specifically, we considered a network in which the probability of connection between pairs was

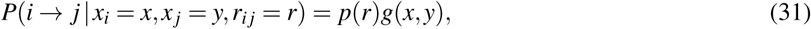
where *p*(*r*) depends on the physical distance *r* between pairs, and the modulator component *g*(*x*,*y*) is not reflection symmetric. This model is itself a two-dimensional Modulator network in which one dimension is physical space, and the other represents a property of the neurons not captured by their spatial location, see Fig. 8A. We assumed that the distribution of distances in samples obtained from the model is close to the sampled distribution in the data (Fig. 8B, left) and that the {*X*_i_}_i_ modulators are independent from neuron to neuron and independent of distances. We assume a Gaussian distribution of the modulator and take *g*(*x*, *y*) to be the weighted sum of the p.d.f. of two bivariate

**Figure 8.**
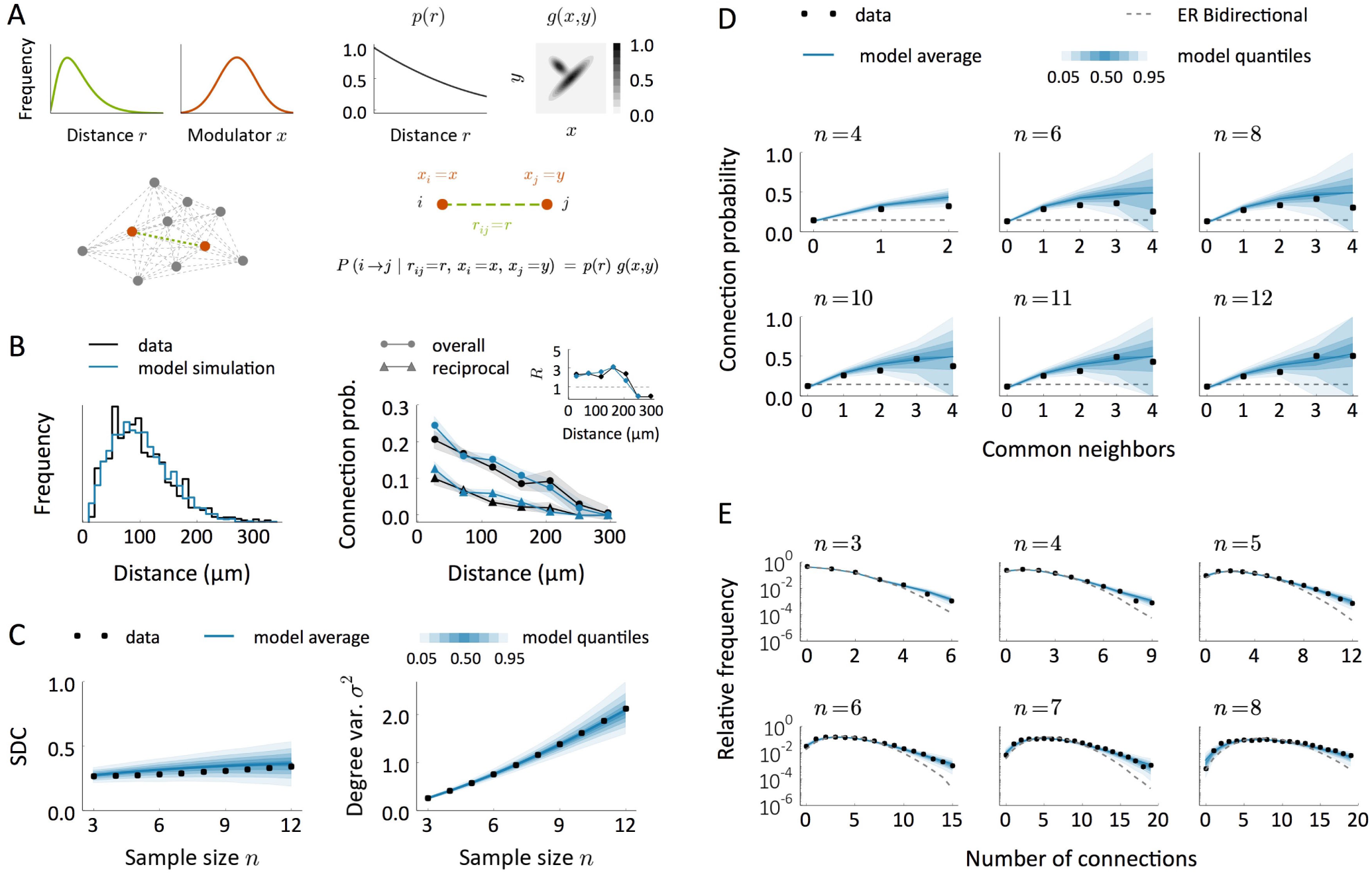
**A** Schematic of a model to explain the observed data. First, neurons are arranged in space so that distances between neuronal pairs follow a given distribution (green). Each neuron has also an associated modulator whose distribution is shown in red. Given a distance-decaying probability *p*(*r*) and a function *g* = *g*(*x*, y), connections are created independently with probability *P(i → j | r_ij_ = r, x_i_ = x, x_j_ = y) = p(r) g(x,y)* **B** Intersomatic distance distribution and connection probabilities as a function of distance in the data (black) and in the model (blue). Inset: number of reciprocal connections relative to random *R* as a function of distance. The model results come from a single replica of the real experiment and shaded regions indicate mean ± SEM. **C** Sample degree correlation SDC and geometric mean of the sample degree variances *σ*^2^ as a function of sample size *n* in the data (black) and in the model (blue). The blue shaded regions indicate quantiles computed from a set of 200 replicas of the real experiment, each performed on an independent network. **D, E** Comparison between model and data in terms of the common neighbor rule (**D**) and the distribution of the total number of connections (E) in samples of size *n*. Dashed lines show the prediction for ER-Bi networks.

Gaussians, one of which breaks the reflection symmetry, see Fig. 8A and Materials and Methods for details. This choice is equivalent to other possible distributions of the modulator as long as *g* is also appropriated transformed. The model successfully captures the observed distance-dependency of the connection probabilities (Fig. 8B right). Note, in particular, that it reproduces the over-representation of reciprocal connections as a function of distance (Fig. 8B right inset). A pure Dis model cannot explain this finding; although the value of *R* evaluated globally would be greater than 1, for any given distance it would be identically 1. Therefore, the increased *R* as a function of distance is a clear signature of additional structure, captured here by our modulator function. The Modulator model also reproduces the sample degree correlation and variance (Fig. 8C), as well as the common neighbor rule and the density of connections in groups of few neurons (Fig. 8D and E).

What is the interpretation of the modulator in this network? The modulator acts as an identifier for each neuron, and neurons with similar modulators will connect in similar ways. Indeed, if the modulator is symmetric we recover a continuous version of a clustered network with heterogeneous membership (Cl-Het). Therefore, the symmetric part of *g*(*x*, *y*) (see plot in Fig. 8A) can be interpreted as clustering: neurons with similar values of *x* are more likely to connect to one-another than to neurons with different values. However, the presence of asymmetry in *g* indicates that connections between clusters are actually hierarchical. Specifically, in our example, neurons with low *x* are likely to connect to similar neurons, and also to neurons with large *x.* On the other hand, neurons with large *x* are likely to connect with similar neurons, but not to neurons with low *x.* In conclusion, the data are consistent with a network in which neurons are connected according to the physical distance between them and their membership in a clustered structure, independent of distance, which itself exhibits hierarchical features.

## Discussion

We have presented three major classes of network models that are compatible with the “non-randomness” reported so far in cortical micro-circuits (Song et al., 2005; Perin et al., 2011). The first is based on a similarity principle: pairs of neurons have associated a notion of distance which modulates the likelihood of the connections between them, in the sense that similar neurons tend to be connected more frequently than different ones. The connections appear independently once the distances between neuronal pairs are known. Distance in this context can represent not only a spatial proximity but any other measure of similarity, for example based on input received from other areas or stimulus selectivity. This family also includes networks where neurons are classified homogeneously into clusters so that connections form preferentially between cells that are in the same cluster. In the second model, neurons are assigned to clusters but there is heterogeneity both in the cluster size and in the number of clusters to which different neurons belong. Connections form with higher likelihood between neurons that coincide in any of the clusters. The third family corresponds to networks where in- and out-degrees of single neurons follow a prescribed joint probability distribution.

Our results show that the three classes of networks can exhibit both an excess of reciprocal connections relative to random and the so-called common neighbor rule for a wide range of parameters. In the case of networks with a specified degree distribution, in- and out-degrees must be positively correlated for the bidirectional connections to be over-represented, meaning that neurons that receive more synapses from the network tend to be the ones that have more outgoing connections, i.e. they are hubs. All of the models can also be similar in terms of the marginal degree distribution in small samples and are in qualitative agreement with previously reported results concerning the number of connections in groups of few neurons. The first important conclusion of our study is therefore that these “non-random” features, rather than being a footprint of a specific topology, seem to arise naturally from several qualitatively distinct types of models.

One of the major difficulties of inferring structural principles from real data is that functional neuronal networks likely encompass thousands of neurons, whereas simultaneous patch-clamp experiments, which provide ground truth for synaptic connectivity, provide samples of only a few neurons at a time. Although the models presented here are based on very different principles, they can be almost indistinguishable from one another given only small sample sizes. Thus, even structures that are distinct globally can exhibit similar properties locally.

A natural question is whether it is possible to define a local measure -i.e., a measure that can be estimated from the study of small samples- that could be used to distinguish between models. We have found such a measure in the *sample degree correlation* (SDC), the correlation coefficient between sample in- and out-degrees. The SDC is, in fact, a particular nonlinear combination of triplet motifs which allows us to correctly classify network models without recourse to training classifiers numerically. Note that a machine learning approach to this problem would require training a classifier on particular instantiations of networks from a given network class; each class encompasses a vast range of possible networks. Therefore training sets would not likely be representative of the class as a whole. A major advantage of our approach, in contrast, is that it allows us to classify networks regardless of the details of every model candidate, which can be difficult to estimate in real situations. For example, in the Distance model the exact shape of the function *p*(*r*) is irrelevant for estimating the SDC, which only depends on the overall connection probability and the over-representation of reciprocal connections. We have also shown that these three model classes are particular cases of a very general model according to which single neurons have an associated property that modulates the connection probability. We call such a property a “modulator”.

We estimated the SDC for distinct data sets from both rat somatosensory cortex and rat visual cortex and found that the structure in those cortical circuits fell outside all three classes of model network in a qualitatively similar way, see Fig.7. Finally, we obtained an excellent fit to the first data set by considering a more general Modulator network in which the probability of connection between neurons depended both on the physical distance between them, as well as on an additional modulator unrelated to distance. In the second data set there is no evidence of distance dependency of connectivity (Song et al., 2005) but the qualitative similarity between data sets in terms of the SDC suggests that a similar non-spatial modulator mechanism might be common to both of them. The structure of this non-spatial modulator could be interpreted as hierarchical clustering, in which connectivity between clusters was asymmetric. However, we cannot rule out that other choices of modulators, which would lead to other interpretations, might provide equally good fits to the data.

The classes of networks that we have explored here are simple enough to be treated analytically. Nature is certainly more complex, and clearly cortical micro-circuits are shaped by other principles, including ongoing synaptic plasticity. We have not considered these mechanisms here. Nevertheless, independent of the mechanisms which shape cortical micro-circuitry, if the topology of the resultant network can be reduced to a modulatory mechanism, then our results show that this modulation involves both a distance dependence and an additional non-spatial component which is asymmetric. This asymmetry means that pre- and post-synaptic components play distinct roles in defining the connection probability.

## Acknowledgments

M.V. has received financial support through the “la Caixa” Fellowship Grant for Post-Graduate Studies, “la Caixa” Banking Foundation, Barcelona, Spain. A.R. acknowledges grants BFU2012-33413 andMTM2015-71509-C2-1-R from the Spanish Ministry of Economics and Competitiveness and grant 2014 SGR 1265 4662 for the Emergent Group “Network Dynamics” from the Generalitat de Catalunya. This work was partially funded by the CERCA program of the Generalitat de Catalunya.

## References

Bastian M, Heymann S, Jacomy M (2009) Gephi: An Open Source Software for Exploring and Manipulating Networks In Third International AAAI Conference on Weblogs and Social Media.

Bock DD, Lee WCA, Kerlin AM, Andermann ML, Hood G, Wetzel AW, Yurgenson S, Soucy ER, Kim HS, Reid RC (2011) Network anatomy and in vivo physiology of visual cortical neurons. Nature 471: 177–182.

Brunel N (2016) Is cortical connectivity optimized for storing information? Nat Neurosci 19: 749–755.

Denk W, Horstmann H (2004) Serial Block-Face Scanning Electron Microscopy to Reconstruct Three-Dimensional Tissue Nanostructure. PLOS Biol 2: e329.

Hill SL, Wang Y, Riachi I, Schürmann F, Markram H (2012) Statistical connectivity provides a sufficient foundation for specific functional connectivity in neocortical neural microcircuits. Proc Natl Acad Sci U S A 109:E2885–E2894.

Hofer SB, Mrsic-Flogel TD, Bonhoeffer T, Hubener M (2009) Experience leaves a lasting structural trace in cortical circuits. Nature 457: 313–317.

Holmgren C, Harkany T, Svennenfors B, Zilberter Y (2003) Pyramidal cell communication within local networks in layer 2/3 of rat neocortex. J Physiol 551: 139–153.

Jiang X, Shen S, Cadwell CR, Berens P, Sinz F, Ecker AS, Patel S, Tolias AS (2015) Principles of connectivity among morphologically defined cell types in adult neocortex. Science 350: aac9462.

Kasthuri N, Hayworth KJ, Berger DR, Schalek RL, Conchello JA, Knowles-Barley S, Lee D, Vázquez-Reina A, Kaynig V, Jones TR, Roberts M, Morgan JL, Tapia JC, Seung HS, Roncal WG, Vogelstein JT, Burns R, Sussman DL, Priebe CE, Pfister H, Lichtman JW (2015) Saturated Reconstruction of a Volume of Neocortex. Cell 162: 648–661.

Kleinfeld D, Bharioke A, Blinder P, Bock DD, Briggman KL, Chklovskii DB, Denk W, Helmstaedter M, Kaufhold JP, Lee WCA, Meyer HS, Micheva KD, Oberlaender M, Prohaska S, Reid RC, Smith SJ, Takemura S, Tsai PS, Sakmann B (2011) Large-Scale Automated Histology in the Pursuit of Connectomes. J. Neurosci. 31: 16125–16138.

Ko H, Hofer SB, Pichler B, Buchanan KA, Sjöström PJ, Mrsic-Flogel TD (2011) Functional specificity of local synaptic connections in neocortical networks. Nature 473: 87–91.

Le Bé JV, Markram H (2006) Spontaneous and evoked synaptic rewiring in the neonatal neocortex. Proc Natl Acad Sci U S A 103: 13214–13219.

Litwin-Kumar A, Doiron B (2014) Formation and maintenance of neuronal assemblies through synaptic plasticity.Nature Communications 5: 5319.

Markram H, Lübke J, Frotscher M, Roth A, Sakmann B (1997) Physiology and anatomy of synaptic connections between thick tufted pyramidal neurones in the developing rat neocortex. J Physiol 500: 409–440.

Markram H, Muller E, Ramaswamy S, Reimann MW, Abdellah M, Sanchez CA, Ailamaki A, Alonso-Nanclares L, Antille N, Arsever S, Kahou GAA, Berger TK, Bilgili A, Buncic N, Chalimourda A, Chindemi G, Courcol JD, Delalondre F, Delattre V, Druckmann S, Dumusc R, Dynes J, Eilemann S, Gal E, Gevaert ME, Ghobril JP, Gidon A, Graham JW, Gupta A, Haenel V, Hay E, Heinis T, Hernando JB, Hines M, Kanari L, Keller D, Kenyon J, Khazen G, Kim Y, King JG, Kisvarday Z, Kumbhar P, Lasserre S, Le Bé JV, Magalhães BRC, Merchán-Pérez A, Meystre J, Morrice BR, Muller J, Muñoz-Céspedes A, Muralidhar S, Muthurasa K, Nachbaur D, Newton TH, Nolte M, Ovcharenko A, Palacios J, Pastor L, Perin R, Ranjan R, Riachi I, Rodríguez JR, Riquelme JL, Rössert C, Sfyrakis K, Shi Y, Shillcock JC, Silberberg G, Silva R, Tauheed F, Telefont M, Toledo-Rodriguez M, Tränkler T, Van Geit W, Díaz JV, Walker R, Wang Y, Zaninetta SM, DeFelipe J, Hill SL, Segev I, Schürmann F (2015) Reconstruction and Simulation of Neocortical Microcircuitry. Cell 163: 456–492.

Mason A, Nicoll A, Stratford K (1991) Synaptic transmission between individual pyramidal neurons of the rat visual cortex in vitro. J. Neurosci. 11: 72–84.

Mountcastle VB (1997) The columnar organization of the neocortex. Brain 120 (Pt 4):701–722.

Nykamp DQ (2007) A mathematical framework for inferring connectivity in probabilistic neuronal networks. Math Biosci 205: 204–251.

Pajevic S, Plenz D (2009) Efficient Network Reconstruction from Dynamical Cascades Identifies Small-World Topology of Neuronal Avalanches. PLOS Comput Biol 5: e1000271.

Perin R, Berger TK, Markram H (2011) A synaptic organizing principle for cortical neuronal groups. Proceedings of the National Academy of Sciences 108: 5419–5424.

Ramaswamy S, Courcol JD, Abdellah M, Adaszewski SR, Antille N, Arsever S, Atenekeng G, Bilgili A, Brukau Y, Chalimourda A, Chindemi G, Delalondre F, Dumusc R, Eilemann S, Gevaert ME, Gleeson P, Graham JW, Hernando JB, Kanari L, Katkov Y, Keller D, King JG, Ranjan R, Reimann MW, Rössert C, Shi Y, Shillcock JC, Telefont M, Van Geit W, Villafranca Diaz J, Walker R, Wang Y, Zaninetta SM, DeFelipe J, Hill SL, Muller J, Segev I, Schürmann F, Muller EB, Markram H (2015) The neocortical microcircuit collaboration portal: a resource for rat somatosensory cortex. Front. Neural Circuits p. 44.

Reimann MW, King JG, Muller EB, Ramaswamy S, Markram H (2015) An algorithm to predict the connectome of neural microcircuits. Front. Comput. Neurosci p. 120.

Roxin A (2011) The Role of Degree Distribution in Shaping the Dynamics in Networks of Sparsely Connected Spiking Neurons. Front Comput Neurosci 5.

Sadovsky AJ, MacLean JN (2013) Scaling of Topologically Similar Functional Modules Defines Mouse Primary Auditory and Somatosensory Microcircuitry. J. Neurosci. 33:14048–14060.

Silberberg G, Gupta A, Markram H (2002) Stereotypy in neocortical microcircuits. Trends Neurosci. 25: 227–230.

Song S, Sjöström PJ, Reigl M, Nelson S, Chklovskii DB (2005) Highly Nonrandom Features of Synaptic Connectivity in Local Cortical Circuits. PLOS Biol 3: e68.

Stetter O, Battaglia D, Soriano J, Geisel T (2012) Model-Free Reconstruction of Excitatory Neuronal Connectivity from Calcium Imaging Signals. PLOS Comput Biol 8: e1002653.

Szentagothai J (1978) The Ferrier Lecture, 1977: The Neuron Network of the Cerebral Cortex: A Functional Interpretation. Proceedings of the Royal Society of London B: Biological Sciences 201: 219–248.

Timme NM, Ito S, Myroshnychenko M, Nigam S, Shimono M, Yeh FC, Hottowy P, Litke AM, Beggs JM (2016) High-Degree Neurons Feed Cortical Computations. PLOS Computational Biology 12: e1004858.

Tomm C, Avermann M, Petersen C, Gerstner W, Vogels TP (2014) Connection-type-specific biases make uniform random network models consistent with cortical recordings. J. Neurophysiol. 112: 1801–1814.

Trachtenberg JT, Chen BE, Knott GW, Feng G, Sanes JR, Welker E, Svoboda K (2002) Long-term in vivo imaging of experience-dependent synaptic plasticity in adult cortex. Nature 420: 788–794.

Wang Y, Markram H, Goodman PH, Berger TK, Ma J, Goldman-Rakic PS (2006) Heterogeneity in the pyramidal network of the medial prefrontal cortex. Nat Neurosci 9: 534–542.

Zuo Y, Yang G, Kwon E, Gan WB (2005) Long-term sensory deprivation prevents dendritic spine loss in primary somatosensory cortex. Nature 436:261–265.

